# Aquaporin-4 mislocalization from astrocyte endfeet prolongs survival in a prion-cerebral amyloid angiopathy model

**DOI:** 10.64898/2026.07.13.732468

**Authors:** Samantha Flores, Amanda Wilpitz, Daniel Ojeda-Juarez, Jin Wang, Garrett Danque, Paige Sumowski, Gail Funk, Adela Malik, Don Pizzo, Emily Richards, Jeffrey J. Iliff, Christina J. Sigurdson

## Abstract

Aquaporin 4 (AQP4) water channels are polarized to astrocytic endfeet at blood vessel interfaces, and lose polarity in vascular diseases, including stroke, chronic traumatic encephalopathy, and Alzheimer’s disease. AQP4 modulates water influx and efflux in the interstitial fluid, yet how AQP4 localization impacts cerebral amyloid angiopathy (CAA) remains poorly understood. Here we show that astrocytic end feet and AQP4 are displaced from amyloid-bearing vessels in a prion-CAA mouse model that expresses GPI-anchorless PrP^C^. Displacing AQP4 genetically through deleting alpha-syntrophin (*Snta1^-/-^*) led to a marked prolongation in survival, together with reduced microglial inflammation and C1q, in prion-CAA-affected mice. Additionally, synaptic structural proteins were better maintained. Finally, the level and distribution of prion aggregates were similar among the mice, indicating that prion conversion and spread was not affected. These results suggest that reducing AQP4 water channel function slows the decline in a vascular amyloid disease by reducing neuroinflammation.

## Introduction

Prion diseases, including sporadic Creutzfeldt-Jakob disease (CJD), are prototypic neurodegenerative disorders induced by PrP^Sc^, an aggregated conformer of the cellular prion protein, PrP^C^ (1). PrP^C^ is glycophosphatidylinositol (GPI)-anchored, however, 10-15% is cleaved from the cell surface by ADAM10 protease and released into the interstitial fluid (ISF). In prion and Alzheimer’s disease, PrP^Sc^ and amyloid-β (Aβ) aggregates, respectively, accumulate and spread extracellularly at the synapse and through the ISF (2), and therapies that slow aggregate spread are limited. PrP^Sc^ and Aβ are partially cleared through ISF-CSF exchange and CSF drainage into venous blood (3, 4), yet aggregates retained in the parenchyma drive metabolic dysregulation (5), spine loss (6, 7), and neuroinflammation (8), promoting neurodegeneration and cognitive decline. Recent reports suggest cerebral ISF flow changes with age (9, 10), which may impair extracellular protein clearance and exacerbate aggregate accumulation (11–13).

CSF-ISF solute exchange and the clearance of protein aggregates and metabolic waste occurs within the glymphatic system, a network of perivascular channels that links CSF influx with the interstitial space of the brain parenchyma (14, 15). Astrocytes serve as a key conduit that regulates water flow from perivascular channels into the interstitial space via AQP4 transmembrane channels (16–18). AQP4 channels are polarized, being concentrated in the astrocytic end feet that ensheath cerebral blood vessels to enable the bidirectional transmembrane flow of water between the vasculature and the ISF (16, 19). On the cytoplasmic side of AQP4, dystrophins and dystrobrevin anchoring proteins form a scaffold with α-syntrophin to maintain AQP4 facing endothelial cells (16, 20). Depleting α-syntrophin disrupts AQP4 polarity, untethering AQP4 from perivascular endfeet and disrupting water influx into the ISF (16, 21). In human AD patients, loss of AQP4 perivascular localization correlates with increased Aβ deposits and clinical AD (22), suggesting a potential link between the AQP4 water channel, CSF-ISF solute exchange, and disease progression.

Astrocyte tonicity regulates the abundance of AQP4 on the cell surface and is impacted by disease states. Vascular injury due to CAA, or acute hypoxia from stroke or trauma, increases AQP4 translocation to the plasma membrane to enhance water permeability, causing astrocytes to swell (cytotoxic edema) (23). Thus, astrocytes change their cellular volume during CNS edema to regulate ISF solute homeostasis (24, 25). Notably, AQP4 knockout mice show reduced brain edema following ischemia, suggesting that AQP4-mediated water influx into astrocytes contributes to cytotoxic edema formation (26). Given that brain edema is a prevalent and therapeutically challenging co-morbidity in neurologic disease, including cancer, stroke, and traumatic brain injury, AQP4’s role in regulating ISF fluid flux has led to its emergence as a potential drug target to mitigate cytotoxic and vascular-derived CNS edema (27, 28).

Although AQP4 deletion is beneficial to mouse models with brain edema, deleting AQP4 in neurodegenerative disease models, including Alzheimer’s disease (AD)-related amyloid-β (APP/PS1 and 5XFAD) and tau mouse models (PS19), exacerbates pathogenic Aβ and phospho-tau pathology, respectively (11–13). Additionally, in human CAA cases, AQP4 immunostaining is enhanced in the brain, particularly at CSF and perivascular interfaces (29). Similarly, in prion disease, AQP4 is upregulated in the brain of CJD patients (30) and experimental prion models (31), yet whether AQP4 expression or localization impacts PrP^Sc^ dissemination or disease progression is largely unknown.

Alterations in ISF transport due to plaques, gliosis, blood brain barrier (BBB) structural defects, or altered astroglial water channels occur in prion disease and may affect disease progression. Here we evaluate the impact of AQP4 localization in the brain of two prion-infected mouse models, WT mice and transgenic mice that express GPI-anchorless (secreted) PrP^C^ [Tg(GPI-PrP)], which develop either cell membrane-bound PrP^Sc^ or prion-CAA, respectively. In the prion-CAA model only, amyloid-bearing vessels showed a dramatic reduction in perivascular astrocytes and AQP4. We crossed α-syntrophin-deficient (*Snta1^-/-^*) mice, which lack perivascular AQP4 localization, to the Tg(GPI^−^PrP) mice, and find that Tg(GPI^−^PrP);*Snta1^-/-^* mice with prion-CAA survive markedly longer than the Tg(GPI^−^PrP) controls. Tg(GPI^−^PrP);*Snta1^-/-^*mice showed less microglial and IFN-linked neuroinflammation and slower prion-CAA-induced neurodegeneration, collectively supporting a model in which displacing AQP4 from perivascular sites reduces prolongs survival. Reducing AQP4 surface expression in early neurovascular disease could be a potent strategy to alleviate certain types of vascular-derived neuroinflammation.

## Results

### Prion cerebral amyloid angiopathy disrupts AQP4 positioning at the blood vessel

We first tested whether AQP4 is displaced from perivascular astrocytic endfeet in the prion-affected brain by investigating prion-infected wild type (WT) mice. Given that certain prion aggregates retain the GPI-anchor (32), we inoculated WT mice with two GPI-anchored prion strains, RML or 22L, or a partially GPI-anchored and anchorless strain, ME7 (33), as well as a mock control (uninfected brain). The brain was collected at terminal disease and immunolabeled for PrP^Sc^, AQP4, and the astrocytic marker GFAP. AQP4 remained in astrocytic end feet irrespective of the presence of PrP^Sc^, with occasional vessels showing more intensely vascular and perivascular AQP4 compared to the control brain (Fig. 1A, Fig. S1). Notably, there was also a diffuse, widely distributed increase in AQP4 labelling in the parenchyma (Fig. 1A), similar to observations reported for human prion-affected brain (30, 34, 35).

**Figure 1.**
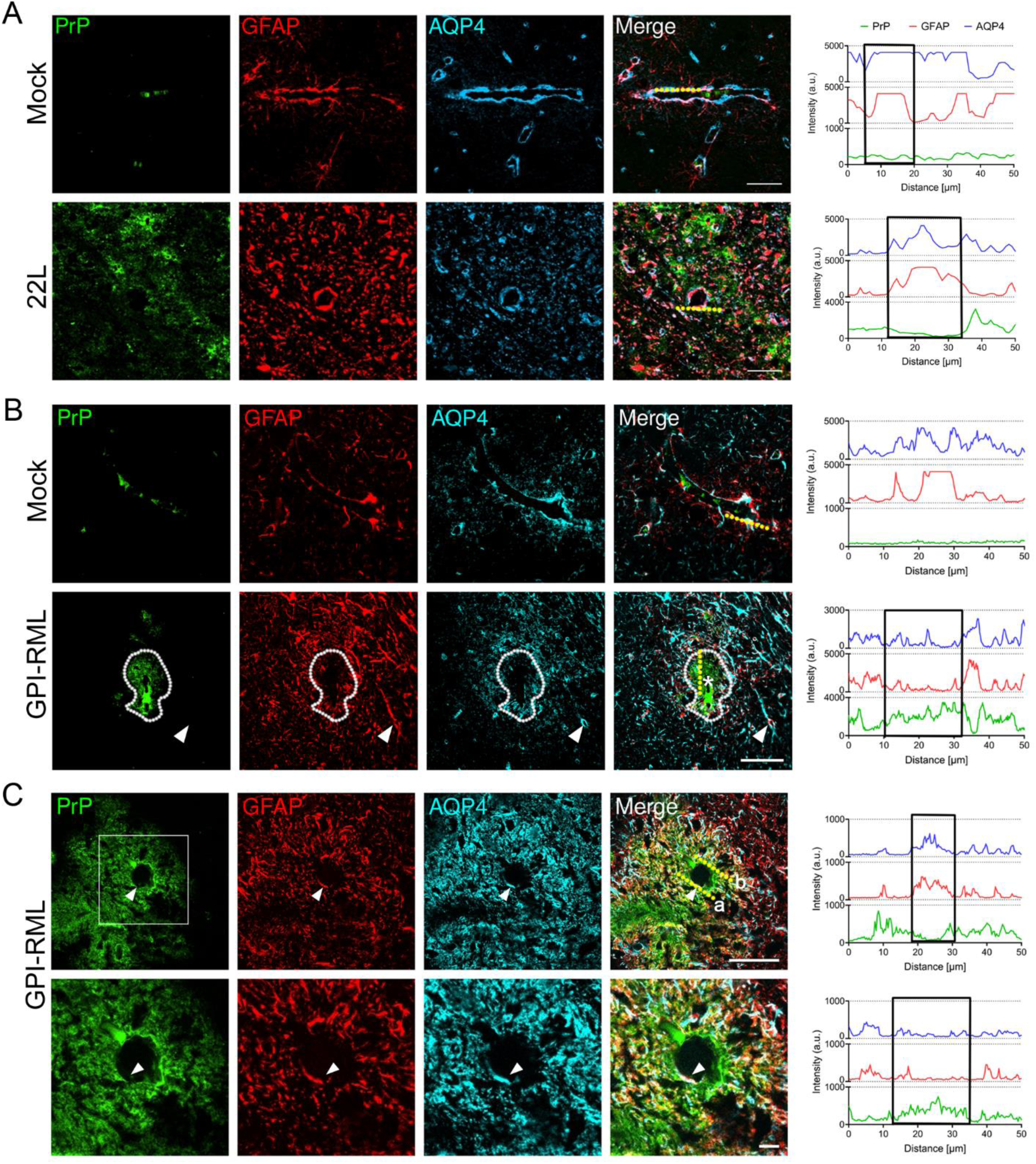
AQP4 redistributes in amyloid-bearing vessels in Tg(GPI-PrP) mice with terminal prion disease. (A) Representative image of thalamus showing mock or prion-affected (22L, GPI-anchored) vessels immunolabeled for PrP^Sc^ (green), GFAP (red), and AQP4 (blue). Traces of each fluorophore from a 50 µm segment of blood vessel (dashed yellow line) show signals of GFAP and AQP4 remain associated with vessels. The black box shows regions of co-localization between GFAP and AQP4. (B) Prion-affected brain (thalamus) (GPI-anchorless RML) shows widespread astrocytosis (red), but a loss of GFAP and AQP4 around a PrP^Sc^-bearing vessel (denoted by *). Arrowhead indicates GFAP and AQP4 localized to a vessel that lacks PrP^Sc^. Traces show that GFAP and AQP4 co-localize in blood vessels in mock brain, but not in GPI-RML-affected brain [note that GFAP and AQP4 are low at sites of abundant PrP^Sc^ (green trace)]. (C) Low (top) and high (bottom) magnification of prion-affected vessels. Shown is a vessel with segments harboring PrP^Sc^ that lack AQP4 (yellow line ‘b’), while segments that lack PrP^Sc^ maintain AQP4 (yellow line ‘a’). The high magnification shows a vessel segment that lacks PrP^Sc^ and maintains GFAP and AQP4 (arrowheads). Scale bars = 50 µm (A, B, and C, upper row), and 10 µm (C, lower row).

We next assessed how GPI-anchorless PrP^Sc^ impacts AQP4 polarity. We used Tg(GPI^−^PrP) mice that accumulate amyloid within and surrounding the vessel wall as a prion-CAA (36). Tg(GPI^−^PrP) mice were inoculated with GPI-anchorless prions (GPI-RML), and brain collected at terminal disease was immunolabeled for PrP^Sc^, AQP4, and GFAP. Perivascular amyloid deposits were encircled by reactive astrocytes (Fig. 1B). Within amyloid-bearing vessels, AQP4 lacked direct association with vessels and was instead abundantly and diffusely distributed beyond the vascular amyloid (Fig. 1C), likely due to displaced astrocytic endfeet (Fig. 1C). Amyloid-free vessels retained perivascular AQP4 similar to the mock-inoculated control brain (Fig. 1B). Notably, in vessels with segmentally distributed amyloid, AQP4 and astrocytic processes redistributed only in the segment of vessel harboring amyloid (Fig. 1C).

### Astrocytic end feet and AQP4 vascular displacement occurs early and is restricted to PrP^Sc^-bearing vessels

To determine the temporal kinetics for AQP4 displacement from vessels, Tg(GPI^−^PrP) mice were inoculated with GPI-RML and the brains were collected at the 40% and 70% timepoints. AQP4 was displaced from vessels harboring amyloid starting at the earliest timepoint, suggesting that AQP4 localization is determined by the presence of amyloid, not by the number of vessels affected or the overall amyloid load (Fig. 2A). At the 40% and 70% timepoints, astrocytic processes retracted only from vessel segments harboring amyloid (Fig. 2B), as observed at terminal disease. Astrocytes were displaced to the amyloid periphery, however, were occasionally embedded within perivascular amyloid with thin processes extending to the vessel wall (Fig. 2B).

**Figure 2.**
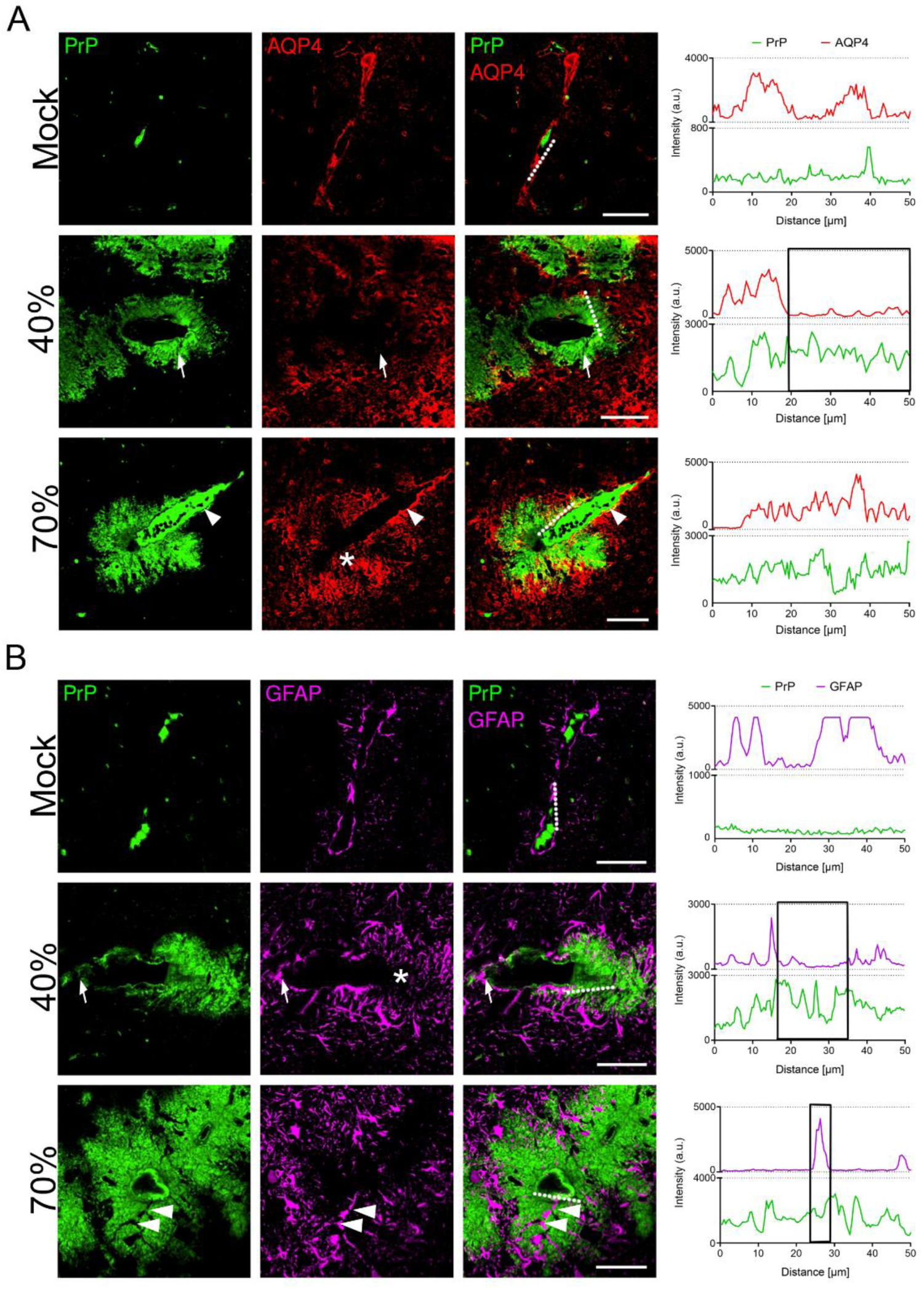
Lack of vascular AQP4 polarization and astrocytic perivascular endfeet in Tg(GPI-PrP) mice, starting in early prion disease (GPI-anchorless RML). (A) At the 40% and 70% timepoints, AQP4 (red) is displaced from vessels harboring PrP^Sc^ (green) (*), with traces showing a lack of co-localization between PrP^Sc^ and AQP4. The 70% timepoint image shows a vessel with PrP^Sc^ within and extending out from one pole. The PrP^Sc^-bearing segment lacks AQP4 (asterisk), while the PrP^Sc^-bearing segment maintains polarized AQP4 (arrowhead). (B) At the 40% timepoint, a vessel lacks astrocytic endfeet (GFAP, magenta), but only at sites that harbor PrP^Sc^ (asterisk). Astrocytic endfeet remain in the PrP^Sc^-free vessel segment (arrows). Traces show little co-localization of the GFAP and PrP^Sc^ spectra (black box). At the 70% timepoint, a vessel harboring abundant PrP^Sc^ shows most astrocytes and processes are peripherally displaced, with a rare astrocyte embedded within the amyloid extends a process to the vessel (arrowheads; black box in traces). Scale bars = 50 µm (A, B, and C).

### Disrupting AQP4 polarity through α-syntrophin depletion (*Snta1^-/-^*) prolongs survival in prion-infected Tg(GPI^−^PrP) mice

AQP4 channels are anchored to the perivascular dystrophin-associated complex by the cytoplasmic, membrane-associated adaptor protein, α-syntrophin (*Snta1*) (16, 17). *Snta1^-/-^*mice express AQP4, but lack the perivascular localization (16). To test how AQP4 distribution impacts survival in mice affected with prion-CAA, Tg(GPI^−^PrP) mice were crossed with *Snta1^-/-^*mice and backcrossed to obtain Tg(GPI^−^PrP);*Snta1^-/-^*mice, hereafter referred to as GPI^−^PrP;*Snta1^-/-^* mice (Fig. 3A). GPI^−^PrP;*Snta1^-/-^* and GPI^−^PrP;*Snta1^+/+^*control mice were inoculated with GPI-anchorless RML prions. The prion-infected GPI^−^PrP mice developed prion disease after 168 ± 7 days. Surprisingly, the GPI^−^PrP;*Snta1^-/-^* mice survived significantly longer (191 ± 5 days) (Fig. 3B), indicating that the AQP4 displacement from astrocytic end feet prolonged survival in mice developing prion-CAA.

**Figure 3.**
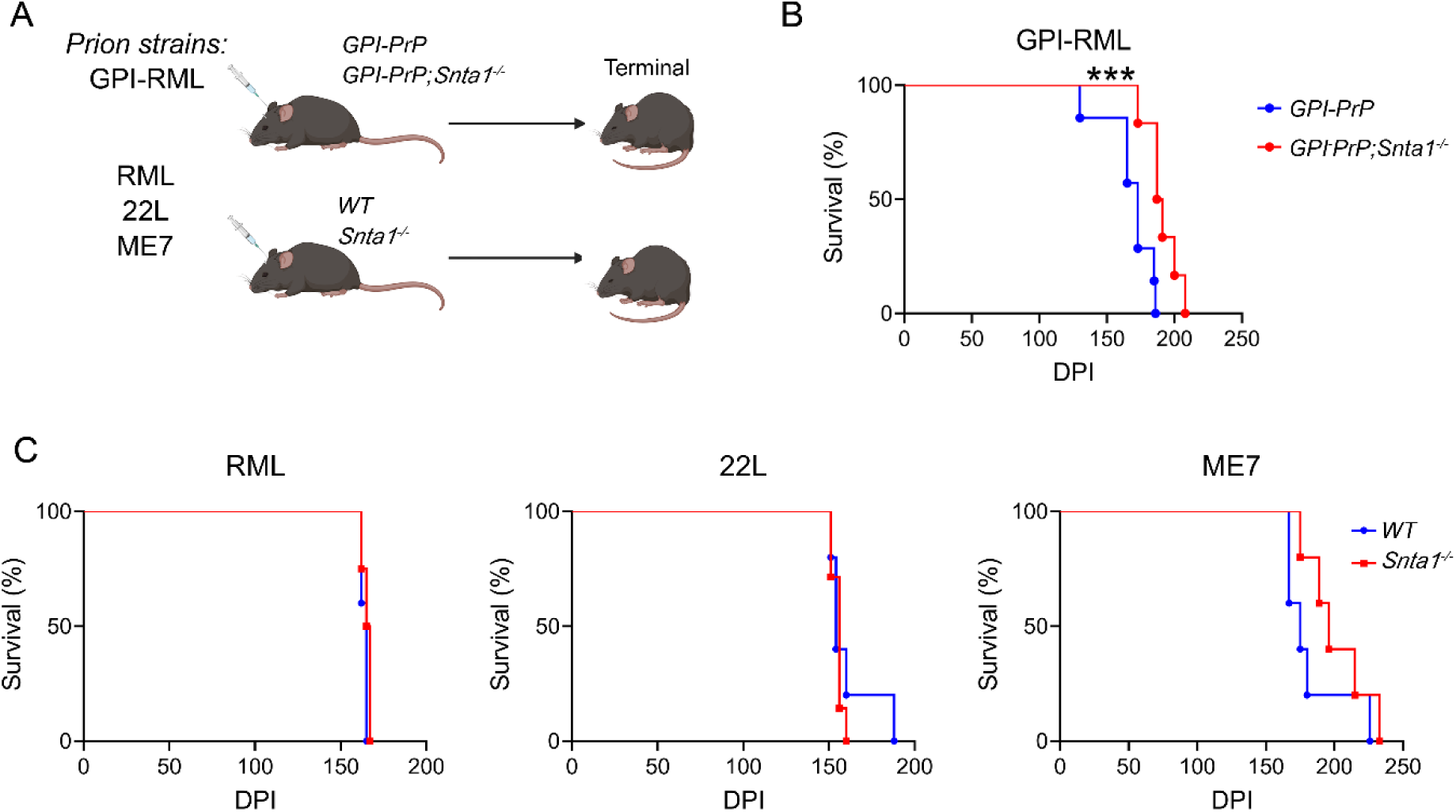
Survival curves of prion-infected *Snta1^-/-^* and control mice. (A) Experimental design of the prion strains inoculated intracerebrally into four mouse lines. (B) Survival curves of the GPI^−^PrP and GPI^−^PrP;*Snta1^-/-^*mice inoculated with GPI-RML prions. (C) Survival curves of the *Snta1^-/-^*and *Snta1^+/+^* mice inoculated with strains RML, 22L, and ME7. *** *P*<0.001, log-rank (Mantel-Cox) test (panels B and C).

GPI-anchored and -anchorless PrP^Sc^ from the strain RML have the same secondary and tertiary structure, yet differ in quaternary structure, as GPI-anchored RML does not form long fibrils (36–38). To assess the impact of α-syntrophin deletion on a GPI-anchored prion infection, *Snta1^-/-^* and *Snta1^+/+^* (WT) mice were inoculated with RML. In contrast to the GPI-anchorless RML-infected mice, RML-infected *Snta1^-/-^* and *Snta1^+/+^* mice showed a nearly identical survival time (168 ± 7 days) (Fig. 3C), indicating that AQP4 displacement had no effect on prion disease progression. To further assess the model for a strain-effect, *Snta1^-/-^* and *Snta1^+/+^*mice were inoculated with two additional prion strains (22L and ME7) (Fig. 3C). Similarly, no differences in survival time were observed, suggesting that AQP4 localization does not impact the disease course when prions are membrane-anchored. Notably, the average survival time was longer for *Snta1^-/-^*-infected with ME7 prions [183 ± 11 days (*Snta1^+/+^*) versus 202 ± 10 (*Snta1^-/-^)* days], however differences were not significant (Fig. 3C).

### Depolarizing AQP4 slows synaptic loss in prion-CAA-affected mice

In AD, *Snta1* gene deletion increases Aβ plaque burden, suggesting that reduced AQP4 localization at perivascular end feet may impair solute clearance, including Aβ oligomers, from the parenchyma (39). To determine how the deletion of AQP4 led to the survival advantage in prion-CAA affected mice, we inoculated an additional cohort of GPI^−^PrP;*Snta1^-/-^* and GPI^−^PrP;*Snta1^+/+^*control mice and collected brain at the same pre-terminal timepoint post-inoculation, at 75% of the incubation period (126 days post-inoculation). We measured PK-resistant PrP^Sc^ levels in the forebrain (cerebral cortex, basal ganglia, thalamus) by western blot and immunohistochemistry. PrP^Sc^ levels and distribution were similar, suggesting that the prolonged survival in the GPI^−^PrP;*Snta1^-/-^* mice was not due to differences in prion conversion kinetics or spread (Fig. 4A-B). The distribution of activated microglia and astrocytes relative to the amyloid deposits also appeared to be similar (Fig. S2). Consistent with no change in PrP^Sc^ load, p62 levels were also unchanged in the GPI^−^PrP;*Snta1^-/-^*mice (Fig. S3), suggesting that the survival benefit was not due to an enhanced clearance of PrP^Sc^ or other proteins.

**Figure 4.**
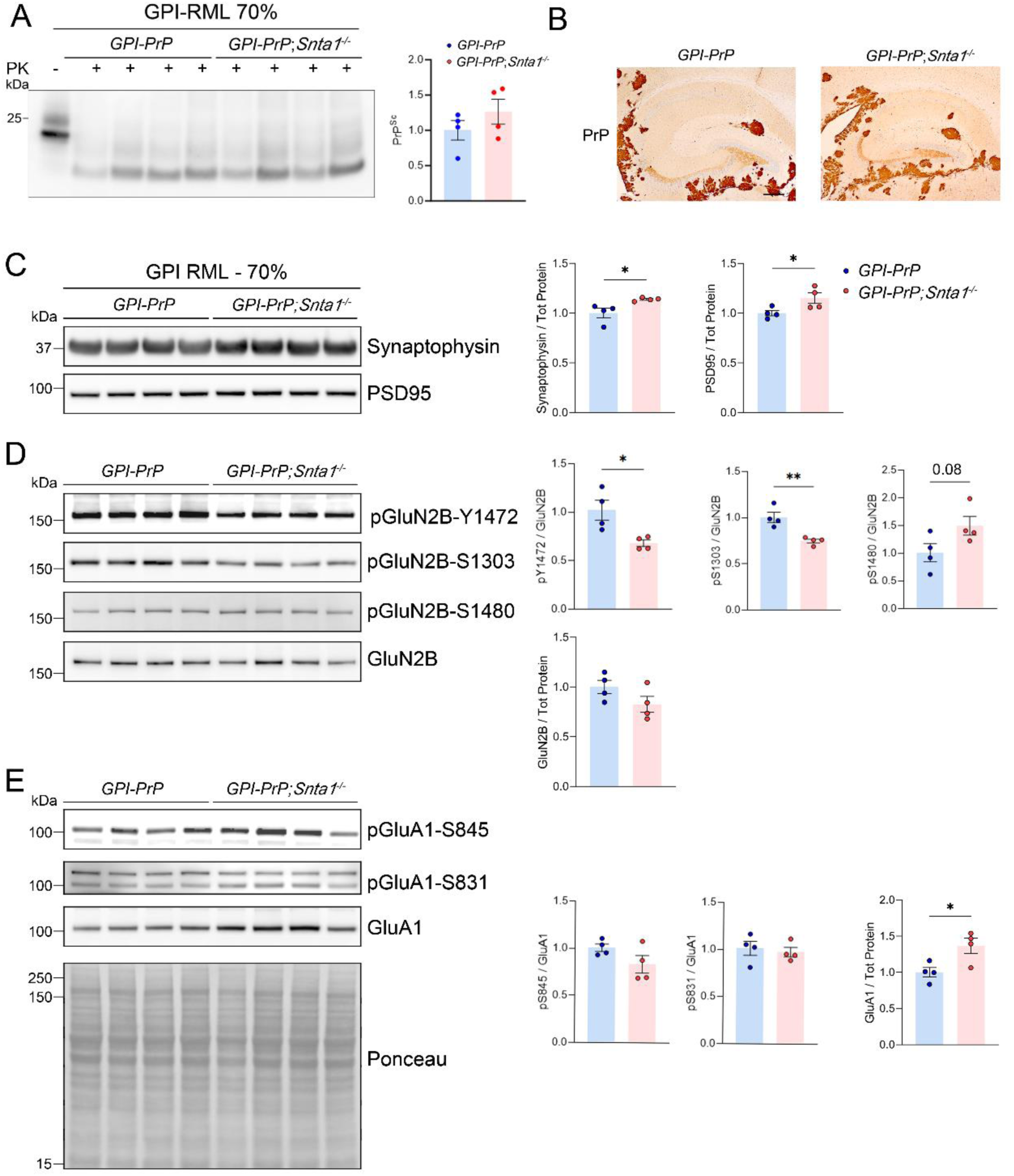
PrP^Sc^ and synaptic proteins in the prion-infected GPI^−^PrP and *Snta1^-/-^*mice at the 70% timepoint. (A) PrP^Sc^ immunoblots and (B) immunolabelled prion-infected GPI^−^PrP and GPI^−^PrP;*Snta1^-/-^*brain reveal similar levels of PK-resistant PrP^Sc^ and distribution of vascular plaques, respectively. Immunoblotting for (C) synaptophysin and PSD95, (D) phosphorylated GluN2B at Y1472, S1303, and S1480, and unmodified GluN2B, and (E) phosphorylated GluA1 at S845 and S831, and unmodified GluA1, together with the quantification. The representative Ponceau stain is from the pGluA1-S845 blot. Bar graphs show mean ± SEM. Unpaired, two-tailed *t*-test, *P < 0.05 and **P < 0.01.

Given the early loss of synapses reported in prion disease (40, 41), we next assessed whether there were measurable structural differences in dendrites or excitatory synapses. We assessed brain lysates for synaptic protein levels, measuring the pre- and post-synaptic proteins, synaptophysin and post-synaptic density protein 95 (PSD95), respectively, by western blot. Strikingly, the prion-infected GPI^−^PrP;*Snta1^-/-^*mice showed more preserved synaptophysin and PSD95 proteins compared to the GPI^−^PrP;*Snta1^+/+^* controls (synaptophysin: 14 +/- 0.01% and PSD95: 15 +/- 0.05% higher) (Fig. 4C), suggesting that synaptic proteins persist longer in mice with AQP4 displacement from the perivascular endfeet.

### The prion-infected GPI^−^PrP;*Snta1^-/-^* brain shows less NMDAR hyperphosphorylation

We previously found that prion-infected mice show increased phosphorylation of NMDA receptors (NMDAR) (GluN2B subunit) and AMPA receptors (AMPAR) (GluA1 subunit) (42, 43), suggestive of alterations in activity-dependent long-term plasticity (44). Therefore, we next measured the levels of phosphorylated glutamate receptors at the 75% timepoint. Notably, prion-infected *Snta1^-/-^* mice showed lower levels of GluN2B phosphorylated at residues Y1472 and S1303, with no change at Y1480, suggesting that displaced AQP4 slows the PrP^Sc^-induced synaptic retention of NMDAR without affecting endocytosis (Fig. 4D). There were no differences in phosphorylated AMPAR (serine 831 or 845), however total GluA1 levels were higher in the *Snta1^-/-^* mice (Fig. 4E). Since the S1303 residue of GluN2B is phosphorylated by CaMKII and protein kinase C (PKC), we used antibodies to measure phosphorylated CaMKII-T286 (a site phosphorylated during increased neuronal activity) and phosphorylated PKC substrates by western blot. There were no differences among the infected mice (Fig. S3), suggesting similar activity of these Ca^2+^-dependent kinases. We also found that cFos levels were similar (Fig. S3).

### Disrupting aquaporin polarization reduces microglial activation, C1q, and alters the inflammatory response to prion-CAA

Astrocytes and microglia are highly activated in prion-infected mice (45). To test whether the *Snta1^-/-^*mice showed altered astrocyte or microglial reactivity, we next measured glial fibrillary acidic protein (GFAP) and ionized calcium-binding adapter molecule 1 (Iba1), respectively, in the GPI-anchorless prion-infected mice by western blot. Notably, there were no differences in astrocytic proteins, GFAP or EAAT2, however Iba1 levels were lower in the GPI^−^PrP;*Snta1^-/-^*mice (Fig. 5A), suggesting less microglial activation. Given microglia phagocytose C1q-tagged synapses and promote synapse loss (46), we next measured C1q. C1q levels were lower in the *Snta1^-/-^* mice (Fig. 5A), consistent with reduced microglial activation and more preserved synaptic proteins. The lower C1q and Iba1, together with the retention of synaptic structural proteins, are consistent with less synaptic tagging, and suggest that AQP4 displacement may be protective due to an attenuated microglial response.

**Figure 5.**
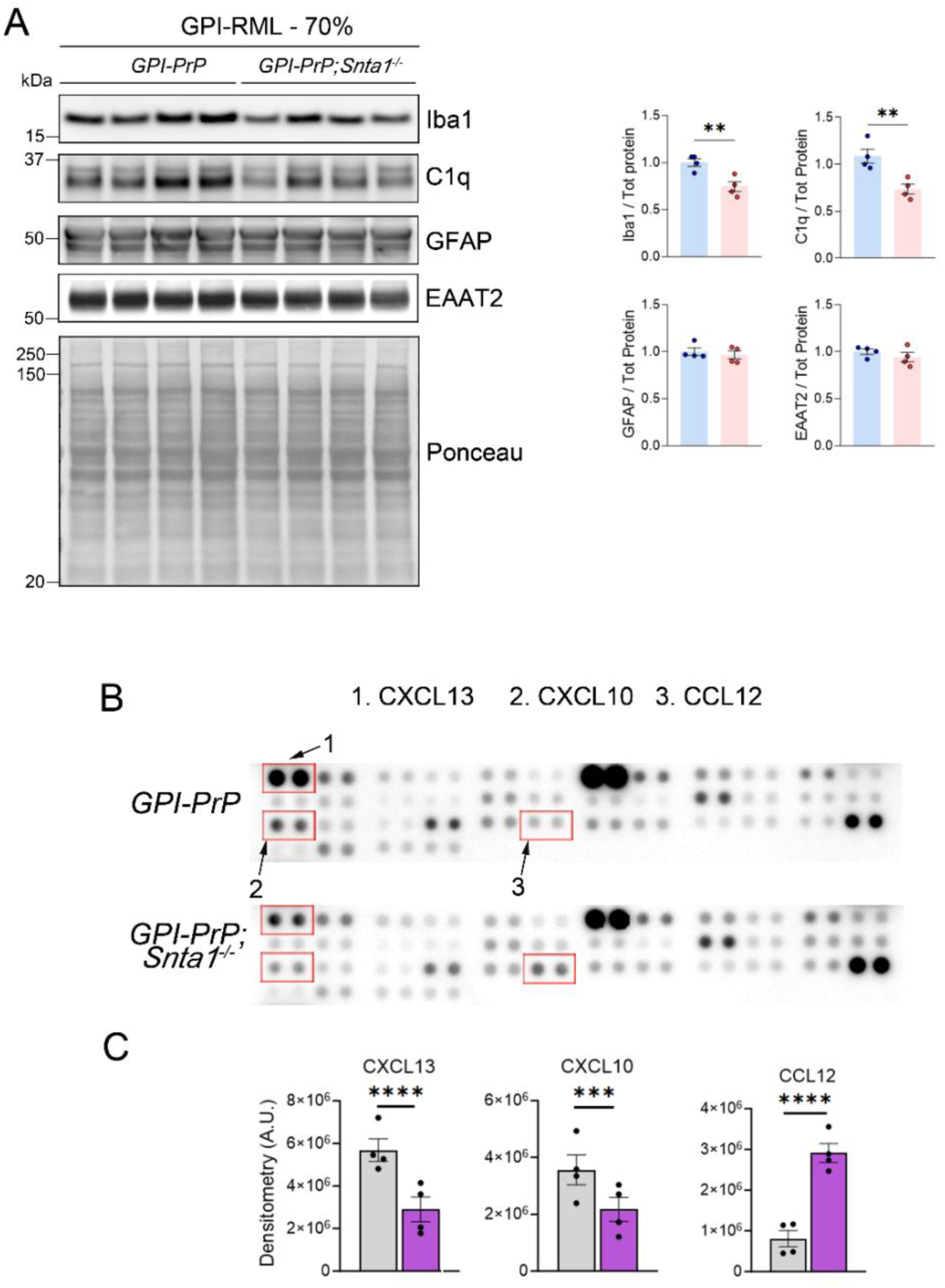
Microglial, complement C1q, and astrocytic proteins in the GPI^−^PrP and GPI^−^PrP;*Snta1^-/-^*brain at the 70% timepoint. (A) Immunoblotting for Iba1, C1q, GFAP, and EAAT2 proteins, and quantification. (B) A representative dot blot assay of cytokines and chemokines from cortical brains extracts shows the cytokine or chemokine level from brain extracts from two individual mice per genotype, normalized by protein level. Red boxes highlight those with significant differences. (C) Quantification below shows results from a total of four mice per genotype. The representative Ponceau stain in (A) is from the Iba1 blot. Bar graphs show mean ± SEM. Unpaired, two-tailed *t*-test, **P < 0.01 (panel A); two-way ANOVA followed by Sidak’s multiple comparisons test (panel C).

To define differences in the neuroinflammatory response associated with the lack of syntrophin, we measured 40 cytokines and chemokines in brain lysates from prion-infected *GPI^−^PrP;Snta1^-/-^*and *Snta1^+/+^* control mice. CXCL10 and CXCL13, which are produced by astrocytes and microglia, were reduced in the *Snta1^-/-^* mice by 39 ± 12% and 49 ± 10%, respectively. CCL12, which is produced by astrocytes and endothelial cells to recruit macrophages, was markedly increased in the *Snta1^-/-^*(259 ± 27%) (Figure 5C-D), together suggestive of a shift in the inflammatory response. In prion disease, CXCL10 and CXCL13 are elevated (47–49) and knockout of their receptor, CXCR3, prolongs survival (50), thus the lower CXCL10 and CXCL13 levels observed in the *Snta1^-/-^* mice may be beneficial.

### Similar synaptic and microglial proteins in mock-inoculated *Snta1^-/-^*and WT mice

To test whether the differences in microglial and synaptic proteins observed in the *Snta1^-/-^* were specific to prion-infected mice, we measured synaptic and glial proteins in mock (uninfected brain)-inoculated mice. We collected *Snta1^-/-^* and WT control mice at 206 days post-inoculation and performed immunoblotting on the cerebral cortex. The astrocytic protein, EAAT2, was modestly higher in the *Snta1^-/-^*group, however, all other proteins measured (synaptophysin, PSD95, GluN2B, GFAP, Iba1) were similar between the *α-Snta1^-/-^* and control mice (Fig. S4). Thus, the differences observed in the prion-infected *α-Snta1^-/-^*mice were specific to the disease state.

### Similar synaptic and microglial proteins in RML-infected *Snta^-/-^*and WT mice

The persistent synapses and lower microglial activation in the GPI^−^PrP;*Snta1^-/-^* mice may not underlie the prolonged survival. Therefore, we compared the levels of synaptic and glial proteins in the RML-infected *Snta1^-/-^* and WT control mice, which did not show a survival difference. There were modest differences in phosphorylated PKC substrates and EAAT2, which were both lower in *Snta1^-/-^*mice (Figs. S5 - S6), however most synaptic and glial proteins were similar between the two groups (Figs. S5 - S6), suggesting that the prolonged preservation of synaptic proteins and the reduced microglial response was specific to the prion-CAA-affected GPI^−^PrP;*Snta1^-/-^*mice.

Collectively, these data indicate that displacing AQP4 in prion-CAA improves survival time. Prion-infected mice with a genetic displacement of AQP4 showed less C1q complement and microglial activation, more preserved synapses, and less NMDAR phosphorylation at a site associated with NMDAR surface retention (1472), which together may underlie the prolonged survival.

## Discussion

AQP4 is a critical water channel in the glymphatic system that facilitates ISF-CSF exchange and metabolic waste clearance, and is increasingly implicated in neurodegenerative disease (51, 52). We found astrocytic endfeet and AQP4 peripherally displaced by amyloid in prion-CAA–affected blood vessels. Increased AQP4 in the cerebral cortex of human sCJD and GSS cases as well as in prion-affected mice has been reported (30, 35), yet the functional impact of AQP4 alterations on prion disease progression has been unclear. Here we demonstrate that AQP4 displacement markedly prolonged survival in prion-infected mice with CAA. Importantly, prion-infected mice with genetically displaced AQP4 (*Snta1^-/-^*) developed less microglial inflammation and showed improved retention of synaptic structural proteins, despite similar PrP^Sc^ levels. Our findings are consistent with a model in which vascular amyloid induces neuroinflammation and complement-mediated synaptic loss that is exacerbated by perivascular astrocytic AQP4.

We propose that vascular amyloid physically displaces astrocytic endfeet from the vessel. Astrocytes showed endfeet retracted from amyloid-laden blood vessels, yet only in vascular segments harboring amyloid, as AQP4 remained perivascular in amyloid-free segments. Moreover, in early disease, some astrocytes were embedded in amyloid with single processes extending to the vessel wall. As vessels accumulated more amyloid with advancing disease, astrocytes and AQP4 were completely displaced, or retracted, from more vascular segments throughout the brain. This displacement and activation of astrocytes may promote perivascular glial scarring in both GPI^−^PrP;*Snta1^-/-^*and GPI^−^PrP mice. However, the prion-infected GPI^−^PrP;*Snta1^-/-^*mice showed a skewed neuroinflammatory profile and reduced synapse loss compared to the GPI^−^PrP mice. By comparison, in AD patients with CAA, astrocyte activation also occurred in the capillary-dominant form of Aβ-CAA, which shows a lower Aβ40:42 ratio and enhanced glial scarring as compared to CAA-affected larger vessels (53, 54). In mouse models, vascular Aβ is associated with a loss of AQP4 polarization, fibrin deposition, and microglial activation (55), consistent with our findings of inflammation within vessels.

The GPI^−^PrP;*Snta1^-/-^* mice with prion-CAA had less C1q, CXCL10, and CXCL13 in the brain, and less microglial activation (Iba1), together with an improved retention of synaptic proteins (synaptophysin and PSD95). These results suggest an altered neuroinflammatory state with reduced complement activation, which may be contributing to preserving synapses and prolonging survival. Specifically, the lower CXCL10 and CXCL13 levels are consistent with less interferon (IFN) signaling, potentially due to altered astrocyte-microglia signaling. In contrast, CCL12 was increased, which has also been reported in intracerebral hemorrhage models to recruit macrophages and T lymphocytes and aggravate injury (56), although the consequences here are not known. Thus, displacing astrocytic AQP4 may be beneficial, potentially by reducing astrocyte swelling, IFN-driven cytokine signaling, and downstream complement-mediated synapse elimination. Whether displacing astrocytic AQP4 reduces IFN-driven cytokine production versus the spread of cytokines from amyloid-bearing vessels remains to be determined.

Consistent with our studies, deleting *AQP4* or *Snta1* improved recovery in models of diffuse neuronal injury, including stroke, bacterial meningitis, and autoimmune encephalitis, potentially by reducing inflammation and edema. Additionally, following intracerebral LPS or stab wound injury, *AQP4*^-/-^ mice showed a reduced cytokine response and less microglial activation (57–60). Interestingly, prion-infected CXCR3 knockout mice reportedly survive longer than controls (50), and CXCL10 is the primary CXCR3 ligand. Our finding of reduced CXCL10 signaling in RML-infected GPI^−^PrP;*Snta1^-/-^* mice suggests that AQP4 depolarization may prolong survival partly through attenuating CXCL10-CXCR3 signaling.

Prior studies indicate that vascular inflammation increases AQP4 expression, astrocytic swelling, and cytokine release, further driving microglial reactivity. Inhibiting HIF1α and AQP4 ameliorated edema in traumatic brain injury (TBI) models that also have vascular injury (61). In contrast, in other diseases, AQP4 may be protective. In an AD model (APP/PS1 mice), AQP4 depletion led to increases in Aβ plaques and exacerbated the loss of synaptic proteins, synaptophysin and PSD95 (11), differing from our findings with prion-CAA in which synaptic proteins were better preserved. Collectively, these findings suggest a context-dependent role of AQP4 in neurovascular and neuroinflammatory disease. Gaining a deeper understanding of how vascular amyloid impacts astrocytic- and microglial-driven inflammation could inform on therapeutic strategies to reduce CNS edema and inflammation in CAA.

We found that the GPI-anchored PrP^Sc^ levels and distribution were unchanged in the *Snta1^-/-^* mice seeded with three prion strains, suggesting that PrP conversion and spread were unaffected by AQP4 depolarization. This finding contrasts with results from experimental AD mice lacking *Snta1*, which show increased Aβ plaques and worsened behavioral deficits (39), implicating AQP4-dependent ISF flow in Aβ peptide clearance. This divergence may reflect differences in how cell membrane-anchored aggregates spread, with GPI-anchored PrP^Sc^ spreading locally to synaptically-linked cells, as evidenced by PrP^Sc^ spread through neuronal circuitry (62) and likely not through the ISF (36). Together these data suggest that the role of AQP4 in ISF flow and particle clearance is more important for diffusible amyloidogenic peptides and oligomers than for membrane-bound aggregates.

In summary, our study demonstrates a critical role for AQP4 localization in the inflammatory response to vascular amyloid. Genetically displacing AQP4 from the astrocytic endfeet along vessels modulated the inflammatory response, slowed the loss of synaptic proteins, and prolonged survival. Given evidence that pharmacologically restoring AQP4 polarization improves outcomes in a TBI model (63), and that the licensed drug trifluoperazine, a calmodulin inhibitor, attenuates edema and neurological deficits following spinal cord injury (23), the current data further support the therapeutic potential of modulating AQP4 levels and distribution, and warrant investigation in vascular dementias, including Aβ-induced CAA.

## Methods

### Mouse and prion inoculations

C57BL/6J, GPI^−^PrP, *Snta1^-/-^*, and GPI^−^PrP;*Snta1^-/-^*mice were housed under specific pathogen-free conditions on a 12-hour light/dark cycle, with ad libitum access to water and standard laboratory chow. At 6 – 8 weeks of age, mice were anesthetized with ketamine and xylazine. Intracerebral inoculations (n = 4 – 7 per group) were performed into the left parietal cortex with 30 μl of 1% 22L, RML, ME7, or GPI-RML prion-infected brain homogenate prepared from terminally ill mice. Clinical terminal stages were identified by the presence of kyphosis, ataxia, hind leg paresis or clasping, and a stiff tail. Additional GPI^−^PrP and GPI^−^PrP;*Snta1^-/-^*cohorts were euthanized at approximated intervals representing 40% and 70% of the disease course.

Upon reaching the clinical endpoint or designated timepoints, brains were harvested and bisected. One hemibrain was snap-frozen on dry ice for biochemical analysis, while the remaining half was fixed in formalin for 2–3 days, treated with 96% formic acid for 1 hour to denature PrP^Sc^, and post-fixed in formalin for histological processing.

### Immunoblotting

Forebrain (cerebral cortex, basal ganglia, thalamus) and isolated cortex were homogenized to a 5% (w/v) concentration in ice-cold RIPA buffer [50 mM Tris HCl (pH 7.4), 150 mM NaCl, 1 mM EDTA] supplemented with phosphatase inhibitors, protease inhibitors (Complete™, Roche), and Benzonase™ (Sigma). Homogenization was performed using a Beadbeater™ (BioSpec). followed by the addition of detergents [1% NP40, 0.5% DOC, 0.1% SDS, and 2% sarcosyl (final)], and incubation for 30-minute on ice. The resulting lysates were cleared by centrifugation at 2000 x g for 10 minutes at 4 °C. Protein concentration in the supernatant was determined via bicinchoninic acid (BCA) assay. Equivalent amounts of protein were reduced with DTT, resolved on 10% Bis-Tris gels, and transferred to nitrocellulose membranes. Membranes were blocked and then incubated with primary antibodies overnight, followed by HRP-conjugated secondary antibodies. Signals were detected using ECL Supersignal West Dura/Femto and captured on a Fuji LAS 4000 imager.

### Enrichment of PrP^Sc^

Sodium phosphotungstic acid (NaPTA) precipitation was performed to concentrate PrP^Sc^ {Wadsworth, 2001 #10696}. Briefly, 5% brain homogenates were mixed with an equal volume of 4% N-lauryl sarcosine and digested with 100 μg/ml proteinase K (PK) for 30 minutes at 37 °C. Following the addition of NaPTA and MgCl_2_, samples were incubated at 37 °C and centrifuged at 18,000 x g for 30 minutes. The resulting pellets were resuspended in 0.1% N-lauryl sarcosine. Samples were resolved by immunoblotting.

### Primary antibodies for western blots

The following antibodies were used for Western blotting: anti-synaptophysin (1:1000, Cell Signaling, 36406S), anti-PSD95 (1:1000, Cell Signaling Technology, 3405T), anti-phosphorylated GluN2B Y1472 (1:1000, Cell Signaling Technology, 4208S), anti-phosphorylated GluN1B S1303 (1:1000, Millipore Sigma-Aldrich, 07-398), anti-phosphorylated GluN2B S1480 (1:1000, PhosphoSolutions, p1516-1480), anti-GluN2B (1:1000, Cell Signaling Technology, 14544), anti-GluN2A (1:1000, Cell Signaling Technology, 4205S), anti-GluN1 (1:1000, Cell Signaling Technology, 5740T), anti-phosphorylated GluA1 S845 (1:1000, Cell Signaling Technology, 8084), anti-phosphorylated GluA1 S831 (1:1000, Cell Signaling Technology, 75574), anti-GluA1 (1:1000, Cell Signaling Technology 13185S), anti-EAAT2 (1:1000, Cell Signaling Technology, 3838S), anti-Iba1 (1:1000, Cell Signaling Technology, 17198), anti-C1q (1:1000, DAKO, A0136), anti-GFAP (1:1000, DAKO, GA52461-2), anti-phosphorylated PKC Substrate (1:1000, Cell Signaling Technology, 2261S), anti-phosphorylated CaMKII (1:1000, Cell Signaling Technology, 12716), anti-CaMKII (1:1000, Cell Signaling Technology 44365), and anti-PrP antibody POM19 (64).

### Immunohistochemistry

Formalin-fixed, paraffin-embedded (FFPE) brain tissues were sectioned at 5 μm and stained with hematoxylin and eosin (HE), or immunostained using antibodies against PrP (SAF84, epitope in the globular domain at the amino acids 160–170 of the mouse PrP), astrocytes (glial fibrillary acidic protein, GFAP), or microglia (Iba1) using antibodies against GFAP (Dako/Agilent Z0334; 1:6000), Iba1 (Wako 019-19741; 1:3000), and PrP^Sc^ (Cayman Chemical; 1:1200). Automated immunolabeling was performed using a Ventana Discovery Ultra system. For PrP^Sc^ immunolabeling, slides were incubated in protease 2 for 20 minutes followed by antigen retrieval in CC1 (Tris-EDTA based; pH 8.5; Ventana) for 64 minutes at 95°C. For GFAP, protease antigen retrieval (P2, Ventana) was used for 16 minutes, and for Iba1, retrieval consisted of CC1 for 40 minutes at 95°C. Primary antibodies were then incubated on the tissue for 32 minutes at 37°C. The secondary antibody (HRP-coupled goat anti-rabbit or anti-mouse; OmniMap system; Ventana) was incubated on the sections for 12 minutes at 37°C. The primary antibody was visualized using DAB as a chromogen followed by hematoxylin as a counterstain. Slides were rinsed, dehydrated through alcohol and xylene and coverslipped.

### Immunofluorescence stain and confocal microscopy

For PrP^Sc^, AQP4, and GFAP triple immunolabelling, tissue sections were stained sequentially first using anti-PrP (SAF84; 1:800) followed by GFAP (Agilent/Dako; 1:4000) and AQP4 antibodies (16473-1-AP; 1:4000) using the tyramide signal amplification system (TSA; ThermoFisher). Slides were stained on a Ventana Discovery Ultra (Ventana Medical Systems, Tucson, AZ, USA). Antigen retrieval was performed using a basic treatment solution (CC1; pH 8.5, Ventana) for 64 minutes at 95 °C followed by a peroxidase quenching step with Discovery Inhibitor (Ventana) for 12 min. Sections then were incubated in anti-PrP antibody for 32 minutes at 37 °C, followed by anti-mouse-HRP (OmniMap Detection Kit, Ventana) and detected using TSA-Alexa 488. The antibodies were stripped from their epitopes by treatment in a citric acid-based solution, pH 6 (CC2, Ventana) for 24 minutes at 95 °C. Subsequently, the slides were incubated with anti-AQP4 antibody for 32 minutes at 37 °C followed by anti-rabbit-HRP (OmniMap Detection Kit, Ventana) and detected using TSA-Alexa 594. Another stripping was done and then the slides were incubated with anti-GFAP antibody and detected with anti-rabbit-HRP with TSA-Alexa647 as the fluorophore. Nuclei were labeled with DAPI, and slides were mounted with fluorescent mounting medium (ProLong^TM^ Gold antifade reagent).

Fluorescence images of AQP4, PrP, and GFAP immunolabeled sections were acquired using the Eclipse Ti2-E (Nikon) microscope in the laser scanning confocal mode (A1R HD, Nikon), and a Nikon Plan Apo ʎ 20x NA 0.75 air objective or Plan Apo ʎ 100x NA 1.45 oil objective. All imaging functions were integrated into the NIS elements software (version 5.42.02: High Content Analysis package).

### Cytokine array

Cytokines were detected using the Proteome Profiler™ mouse cytokine array kit, panel A (R&D systems, ARY006) following manufacturer’s instructions with minor modifications. Briefly, protein concentrations were determined for GPI^−^PrP and GPI^−^PrP;*Snta1^-/-^* brain homogenates in PBS by BCA. Proteins (300 µg) were diluted to 1 ml in lysis buffer (lysis buffer 6) with protease inhibitors (Complete™). Membranes were incubated with protein overnight at 4 °C, followed by incubation with detection antibody cocktails, and streptavidin-HRP for detection. The immunoblots were developed using a chemiluminescent substrate (ECL Supersignal West Femto, ThermoFisher Scientific) and visualized on a Fuji LAS 4000 imager. Densitometry analysis was performed using MultiGauge software (Fujifilm).

### Statistical analysis

All experimental data were analyzed using GraphPad Prism software. For comparisons between two specific groups, an unpaired, two-tailed Student’s *t*-test. Survival data and the progression of clinical disease stages were evaluated using Kaplan-Meier survival curves, with statistical significance determined by the log-rank (Mantel-Cox) test. For the cytokine array quantification, a two-way ANOVA followed by Sidak’s multiple comparisons test was performed. A *P*-value of less than 0.05 was considered statistically significant.

## Acknowledgments

Microscopy and image analysis were performed at the Nikon Imaging Center at UC San Diego. We thank the animal care staff at UC San Diego for excellent animal care.

