## Supplemental Figure 1 - 7 for "Aquaporin-4 mislocalization from astrocyte endfeet prolongs survival in a prion-cerebral amyloid angiopathy model"

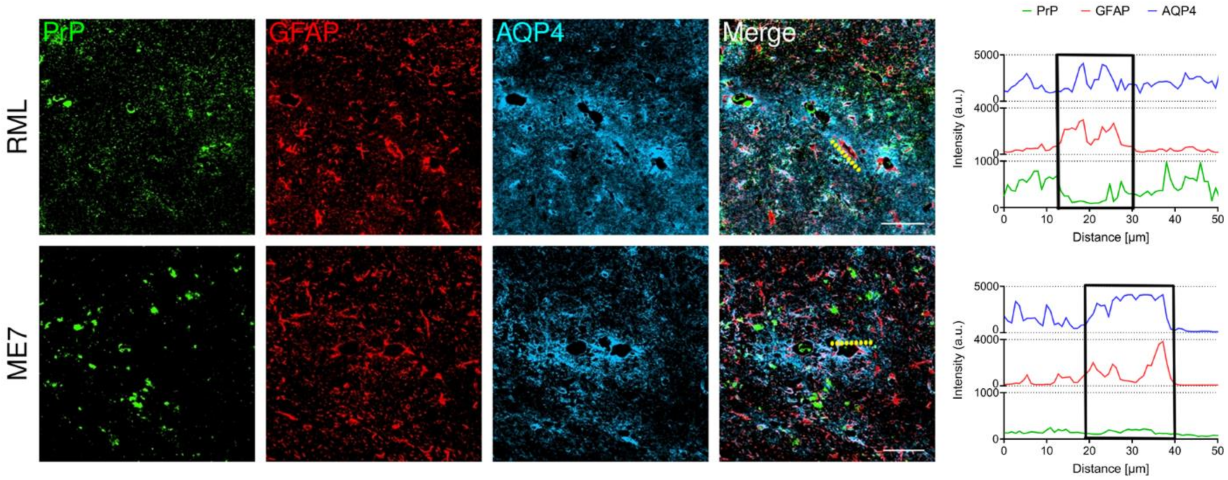


**Supplementary figure 1.** AQP4 is maintained in perivascular astrocytic endfeet in GPI-anchored prion (RML and ME7)-affected brain and is abundant around vessels. (A) Representative image of prion-affected brain immunolabeled for $PrP^{Sc}$ (green), GFAP (red), and AQP4 (blue). Traces show the signal of a 50 µm segment of blood vessel (yellow line) with vessel-associated GFAP and AQP4, but not $PrP^{Sc}$. Black box shows co-localization between GFAP and AQP4. RML: hippocampus and ME7: thalamus. Scale bar = 50 µm

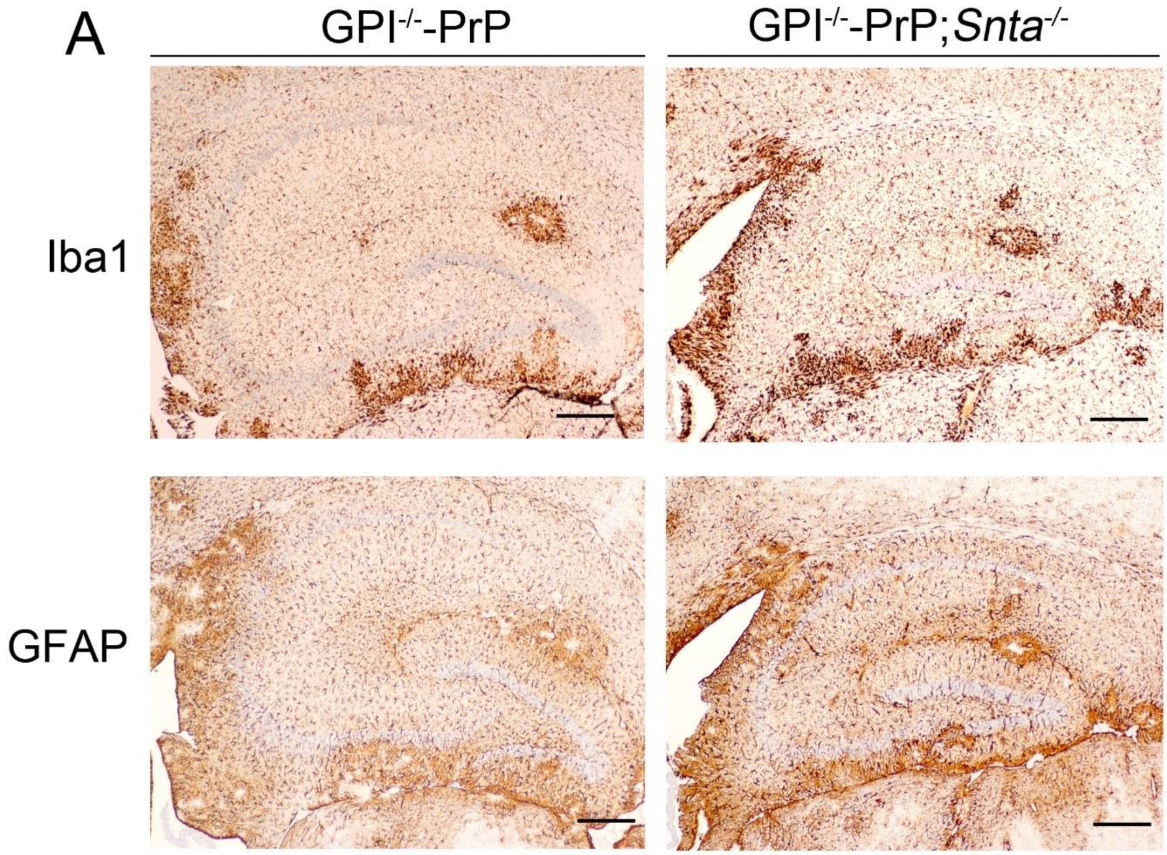


**Supplementary figure 2.** Brain at hippocampus from prion-infected GPI$^{-}$PrP and GPI$^{-}$PrP;*Snta1*$^{-/-}$ mice immunolabeled for microglia (Iba1) and astrocytes (GFAP). Scale bar = 500 µm.

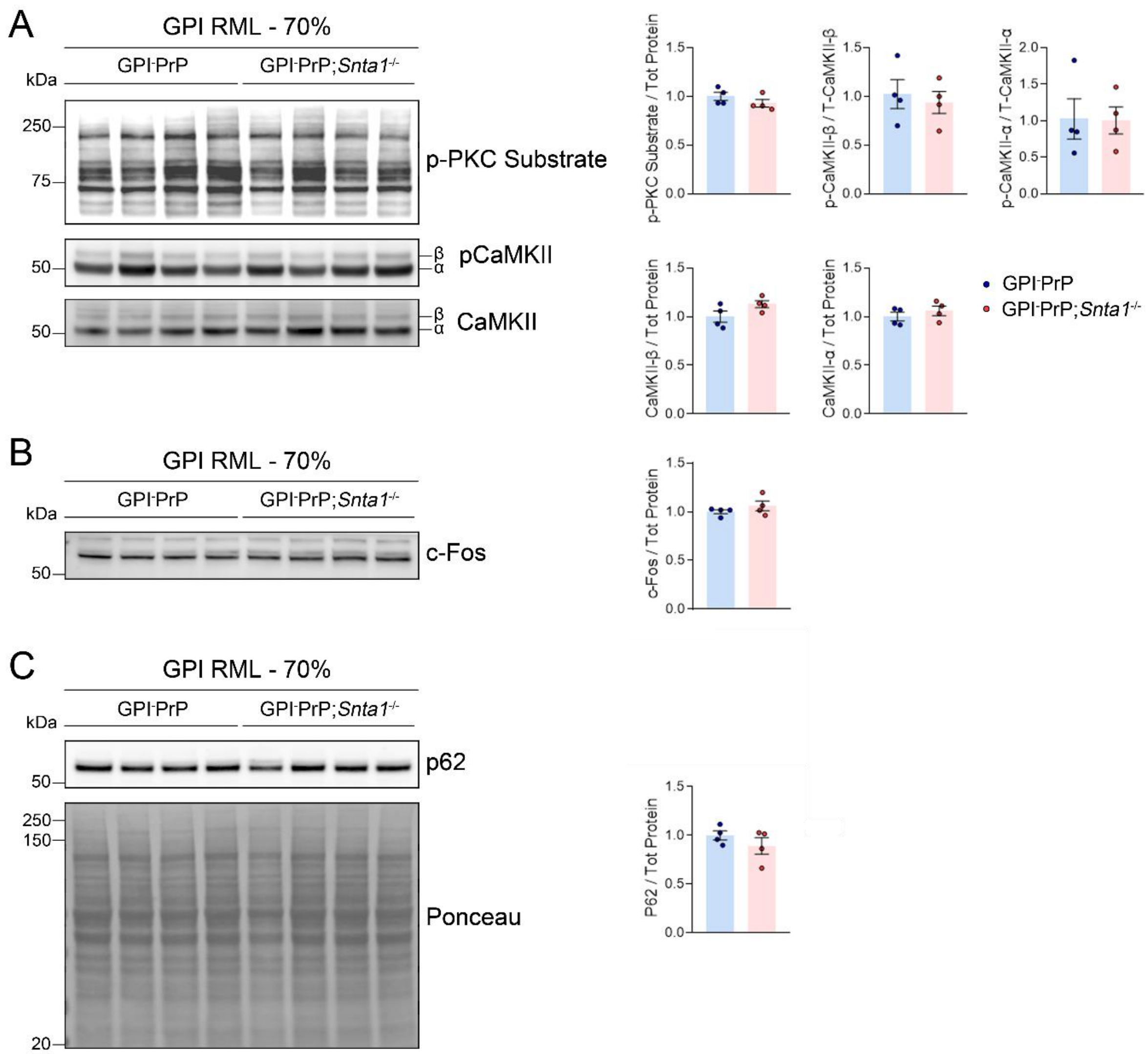


**Supplementary figure 3.** Quantification of GPI$^-$PrP and GPI$^-$PrP;*Snta1*$^{-/-}$ brain for active kinases, neuronal activity, and autophagy-related proteins in the cerebral cortex at the 70% timepoint. Immunoblotting for (A) phosphorylated PKC substrates and CaMKII, (B) cFos, and (C) p62, together with the quantification. The representative Ponceau stain is from the p62 blot. Unpaired, two-tailed *t*-test.

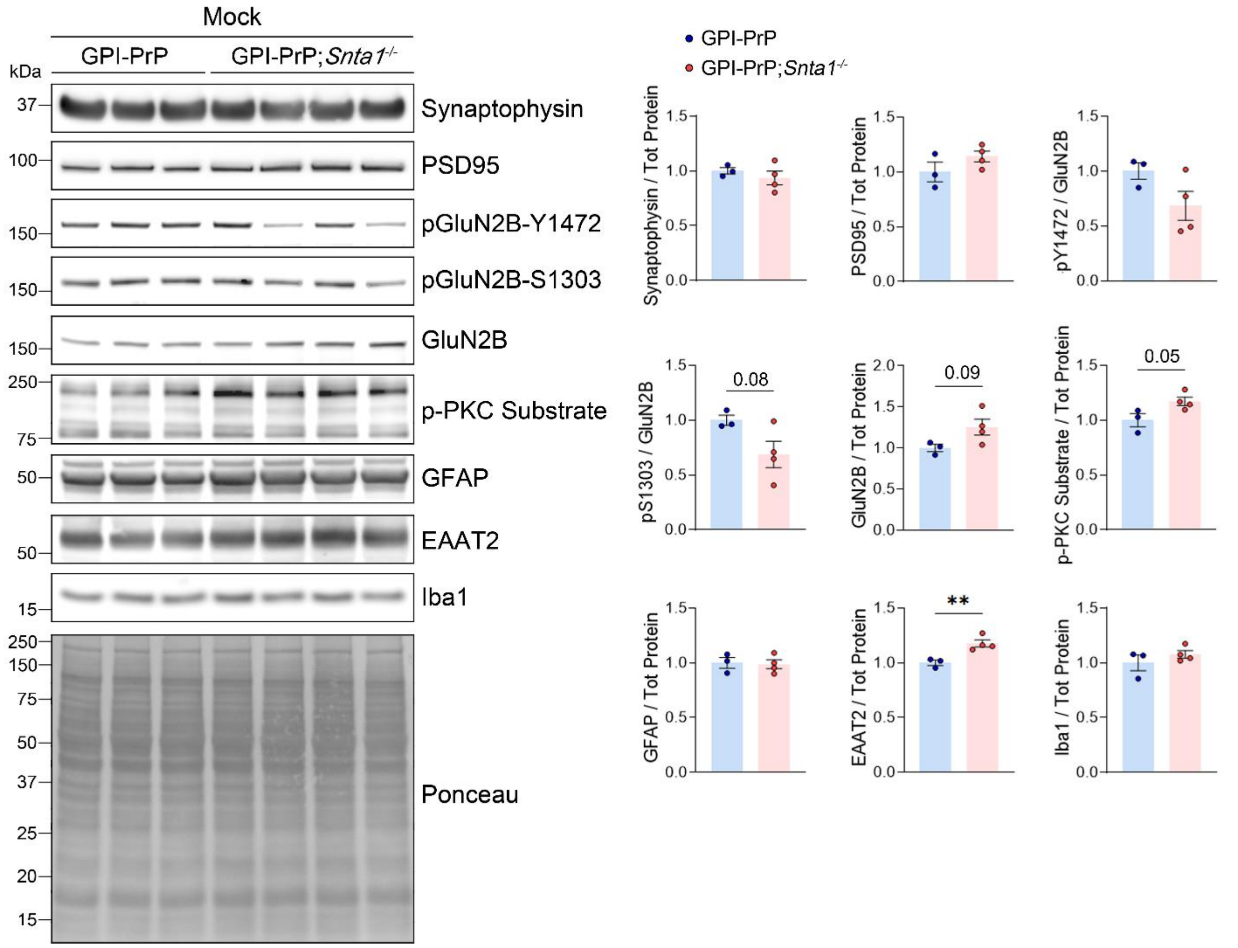


**Supplementary figure 4.** Quantification of synaptic proteins and glial proteins in mock-inoculated *Snta1*$^{+/+}$ and *Snta1*$^{-/-}$ brain. Immunoblotting for synaptophysin and PSD95, phosphorylated GluN2B at Y1472, S1303, and GluN2B, phosphorylated PKC substrates, as well as astrocytic proteins GFAP, EAAT2, and Iba1 together with their quantification. The representative Ponceau stain is from the Iba1 blot. Unpaired, two-tailed *t*-test, $^{**}P < 0.01$.

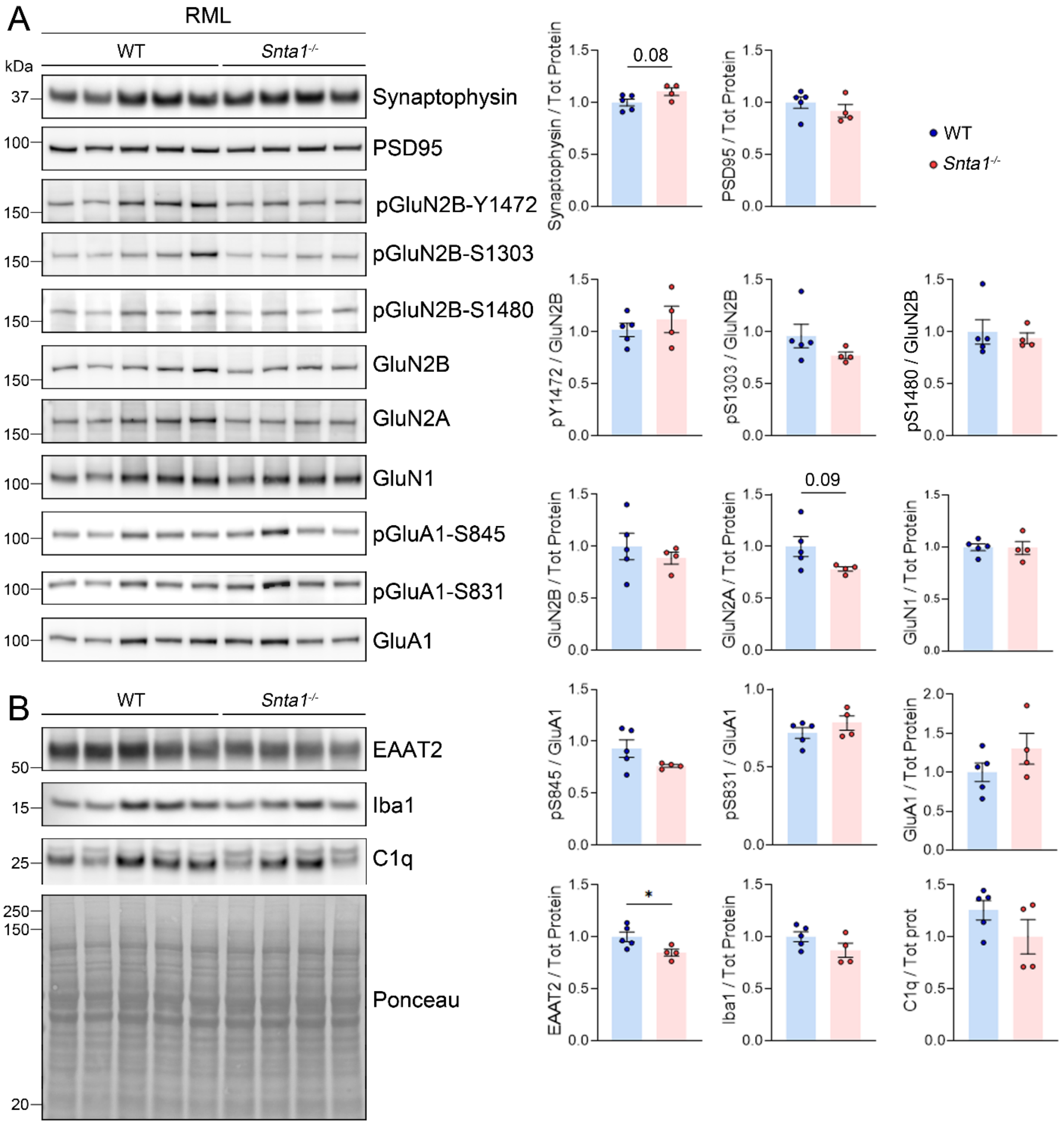


**Supplementary figure 5.** Quantification of synaptic and glial proteins in RML-infected *Snta1*$^{+/+}$ and *Snta1*$^{-/-}$ brain. Immunoblotting of brain lysate for (A) synaptophysin, PSD95, phosphorylated glutamate receptors, and (B) glial proteins, together with their quantification. The representative Ponceau stain is from the C1q blot. Unpaired, two-tailed *t*-test, *P < 0.05.

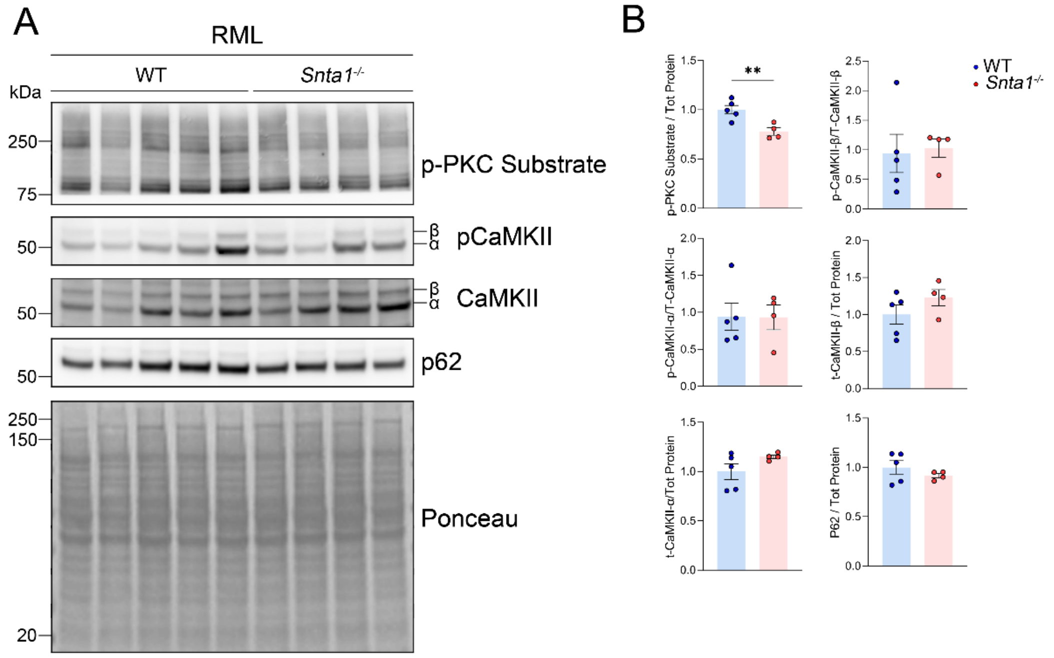


**Supplementary figure 6.** Quantification of kinases and p62 in RML-infected *Snta1*$^{+/+}$ and *Snta1*$^{-/-}$ brain. Immunoblotting of brain lysate for phosphorylated PKC and CaMKII, as well as p62, together with their quantification. The representative Ponceau stain is from the p62 blot. Unpaired, two-tailed *t*-test, **P < 0.01.

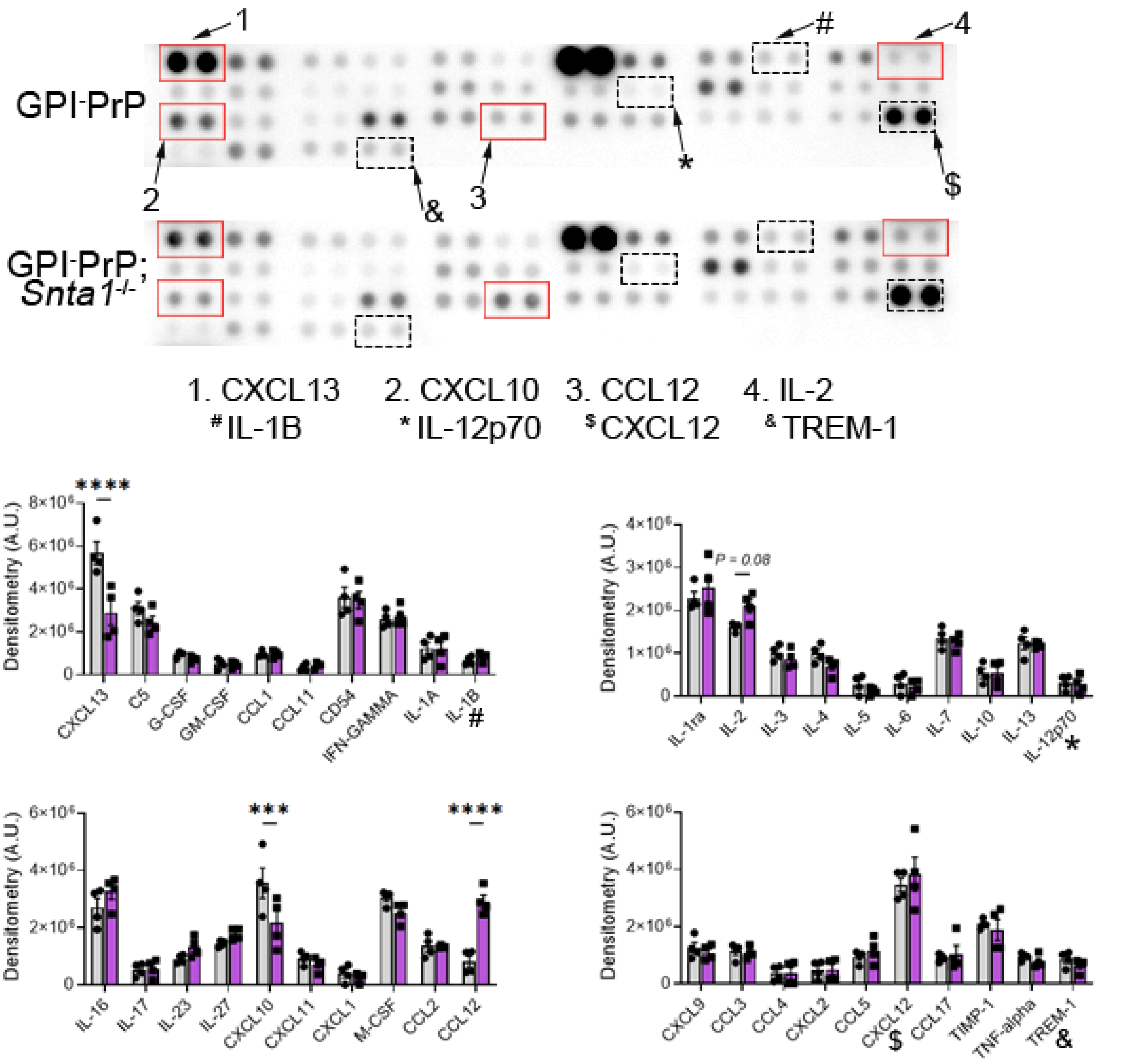


**Supplementary figure 7.** Cytokines and chemokines in cortical brains extracts from GPI⁻PrP and GPI⁻PrP;*Snta1*$^{-/-}$ brain at the 70% timepoint. Representative dot blot assay showing the cytokine or chemokine level from brain extracts from a GPI-PrP and GPI-PrP;*Snta*$^{-/-}$ mouse, normalized for protein level. Red boxes highlight proteins with significant differences. Quantification below shows a total of four mice per genotype. The order of cytokines presented in the graphs sequentially matches their left-to-right spatial arrangement on the dot blot arrays. Symbols (#, *, $, and &) serve as visual reference markers to aid in aligning specific cytokines between the blots and the graphs.
